# Reconstructing NOD-like receptor alleles with high internal conservation in *Podospora anserina* using long-read sequencing

**DOI:** 10.1101/2025.01.13.632504

**Authors:** S. Lorena Ament-Velásquez, Brendan Furneaux, Sonia Dheur, Alexandra Granger-Farbos, Rike Stelkens, Hanna Johannesson, Sven J. Saupe

## Abstract

NOD-like receptors (NLRs) are intracellular immune receptors that detect pathogen-associated cues and trigger defense mechanisms, including regulated cell death. In filamentous fungi, some NLRs mediate heterokaryon incompatibility, a self/non-self recognition process that prevents the vegetative fusion of genetically distinct individuals, reducing the risk of parasitism. The *het-d* and *het-e* NLRs in *Podospora anserina* are highly polymorphic incompatibility genes (*het* genes) whose products recognize different alleles of the *het-c* gene via a sensor domain composed of WD40 repeats. These repeats display unusually high sequence identity maintained by concerted evolution. However, some sites within individual repeats are hypervariable and under diversifying selection. Despite extensive genetic studies, inconsistencies in the reported WD40 domain sequence have hindered functional and evolutionary analyses. Here we demonstrate that the WD40 domain can be accurately reconstructed from long-read sequencing (Oxford Nanopore and PacBio) data, but not from Illumina-based assemblies. Functional alleles are usually formed by 11 highly conserved repeats, with different repeat combinations underlying the same phenotypic *het-d* and *het-e* incompatibility reactions. Protein structure models suggest that their WD40 domain folds into two 7-blade β-propellers composed of the highly conserved repeats, as well as three cryptic divergent repeats at the C-terminus. We additionally show that one particular *het-e* allele does not have an incompatibility reaction with common *het-c* alleles, despite being 11-repeats long. Our findings provide a robust foundation for future research into the molecular mechanisms and evolutionary dynamics of *het* NLRs, while also highlighting both the fragility and the flexibility of *β*-propellers as immune sensor domains.

## Introduction

NOD-like receptors (NLRs) are a class of almost universally conserved intracellular immune receptors that play crucial roles in animal, plant, fungal, and bacterial host defense systems (Jones et al. 2016; Urbach and Ausubel 2017; Dyrka et al. 2014; Kibby et al. 2023). Sometimes referred to as cellular “guardians”, NLRs can sense cues of the unwanted invasion of nonself entities, such as pathogen-derived molecules or pathogen-induced modifications of host cells (Duxbury et al. 2021). Typically, NLRs have a tripartite domain architecture and function through ligand-induced oligomerization (Fu et al. 2024; Gao et al. 2022; Hu and Chai 2023). When a non-self ligand binds to the C-terminal domain (the “sensor”), which is composed of superstructure-forming repeats, it triggers the multimerization of the central nucleotide-binding and oligomerization domain (NBD). This change in the NBD, in turn, activates the N-terminal effector domain that usually leads to regulated cell death (Maekawa et al. 2023). Given this general mode of action, the sensor domain can be under strong selective pressure to keep up with the evolution of pathogens, which change constantly to avoid detection (Kibby et al. 2023; Allen et al. 2004; Melepat et al. 2024).

Filamentous fungi possess large and diverse repertoires of NLRs (Daskalov et al. 2020; Dyrka et al. 2014; Wojciechowski et al. 2022). However, only a few have been functionally characterized, all within the context of heterokaryon or vegetative incompatibility — a self/non-self recognition mechanism occurring between strains of the same species (Daskalov 2023). Growth in filamentous fungi is accomplished by extending their cells or hyphae, by branching, and by fusing with other cells, leading to the possibility of fusing with other individuals (Glass and Dementhon 2006; Harris 2006). This vegetative fusion with non-self poses a great risk, since it opens the door for intracellular parasites such as mycoviruses and selfish organelles, including nuclei (Bastiaans et al. 2016; Debets et al. 2012; Debets and Griffiths 1998; Biella et al. 2002). As a form of defense, different individuals can fuse successfully only if they are compatible at a set of specific loci, termed heterokaryon incompatibility (*het*) genes, some of which are NLRs. Mirroring the innate immune response of other eukaryotes and bacteria, the *het* NLRs trigger regulated cell death of the fused incompatible hyphae, preventing the exchange of cytoplasm and hence parasites (Gonçalves et al. 2017). In plate cultures, this phenomenon can be observed as a line of dead cells in the contact zone between two incompatible strains, called the “barrage” (Esser 2016). In accordance with their self/non-self recognition function, *het* genes in general are highly polymorphic at the population level, displaying signatures of balancing selection (Auxier et al. 2024; Milgroom et al. 2018; Wu et al. 1998).

Among filamentous fungi, *Podospora anserina* has one of the best-studied repertoires of *het* genes (Esser 2016; Pinan-Lucarré et al. 2007). Early classical genetics work on a collection of 16 strains collected in France determined the existence of nine *het* loci (Bernet 1967; Rizet and Esser 1953), all of which have now been cloned (reviewed in Clavé et al. 2024). From these genes, *het-r*, *het-d*, and *het-e* are paralogs from the same NLR type, collectively known as HNWD genes based on their domain architecture (Paoletti et al. 2007). Specifically, HNWD genes are characterized by having a TIR-related <u>H</u>ET effector domain at the N-terminus, an NBD of the <u>N</u>ACHT type, and a sensing domain formed by <u>WD</u>40 repeats at the C-terminus (**Figure 1A**). WD40 domains in general form doughnut-like (toroidal) folds called β-propellers assembled from six to eight repeats (Fülöp and Jones 1999). While many NLRs have WD40 domains, the HWND sensor domain is peculiar in several aspects. On the one hand, the individual WD40 repeats display high sequence identity, ranging from over 80% to 100% within each gene (Saupe et al. 1995a; Espagne et al. 2002; Paoletti et al. 2007). It is proposed that such level of high internal conservation (HIC) reflects concerted evolution of the repeats through unequal crossing-overs or other recombination events that cause high mutation rates (Chevanne et al. 2010; Paoletti et al. 2007; Saupe 2000). This process can add or remove repeat units, leading to length polymorphism in natural populations. On the other hand, while being overall very similar, the individual repeats also show extensive variability at four specific codon positions under diversifying selection, which map to amino acid residues predicted at the interaction surface of the β-propeller (Paoletti et al. 2007).

**Figure 1.**
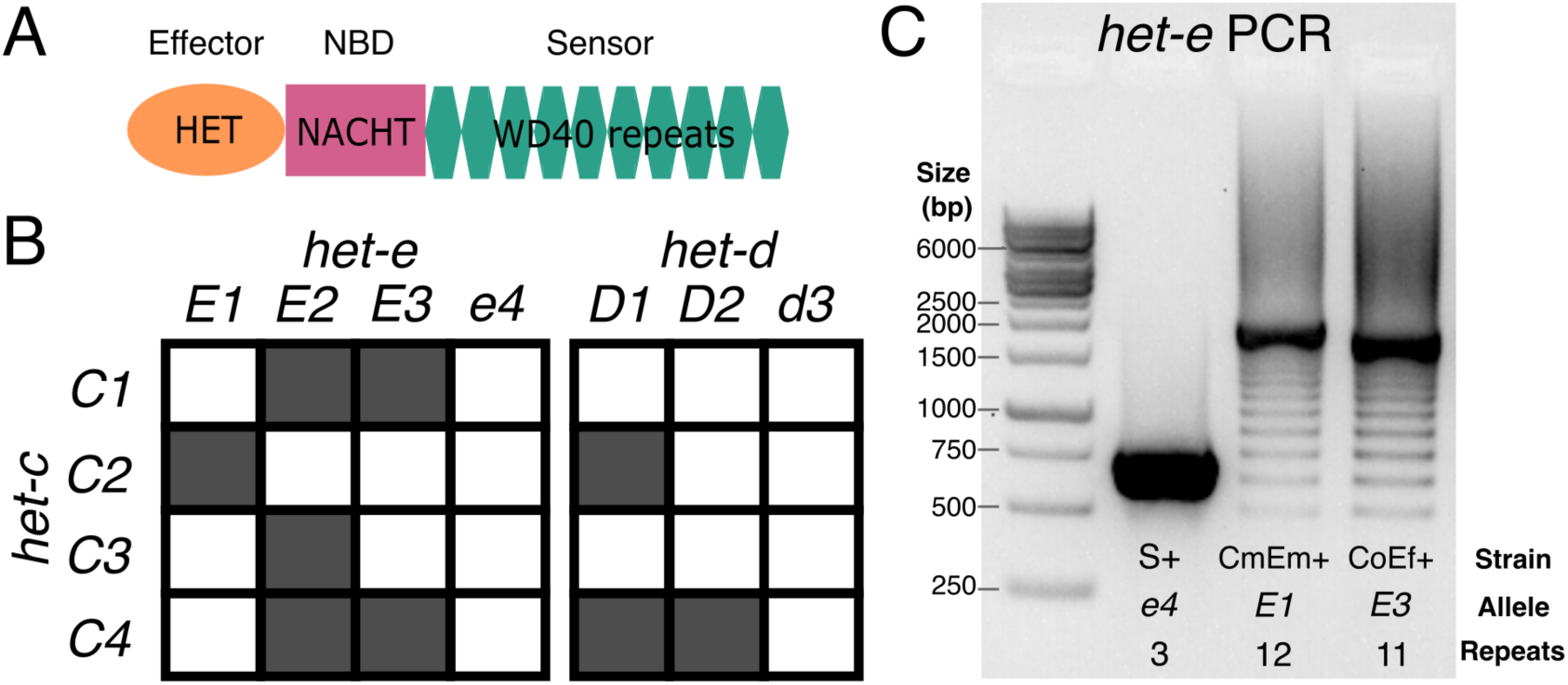
Primer on the *het-d* and *het-e* genes. (A) The domain structure of an HNWD NLR. (B) Incompatibility interactions between the most common *het-c* alleles and those of *het-e* and *het-d*. Shaded squares indicate a vegetative incompatibility reaction, while white squares indicate compatibility, following Saupe et al. (1995b). (C) A typical PCR result when amplifying the WD40 domain of an HNWD gene from genomic DNA, in this case *het-e* (1% agarose gel). Three strains with known *het-e* alleles are shown. NBD: nucleotide-binding domain.

Previous research has shown that the number and sequence of WD40 repeats determine the allele specificity of a given HNWD paralog (Chevanne et al. 2010; Daskalov et al. 2015; Espagne et al. 1997; Saupe et al. 1995a). For example, the product of one specific *het-r* allele of 11 repeats (also known as *het-R* or just *R*) recognizes one allele of the *het-v* locus, triggering the vegetative incompatibility reaction (Chevanne et al. 2010). Other combinations and numbers of repeats are not reactive (known as *r*). Meanwhile, the products of the *het-d* and *het-e* genes recognize the same target, the glycolipid transfer protein coded by the *het-c* gene (Bernet 1967; Saupe et al. 1995b). The *het-c* gene itself is polymorphic and its different (phenotypic) alleles can be defined by their interaction with *het-d* and *het-e* (Bernet 1967). For instance, the *C2* allele triggers an incompatibility reaction with a specific *het-e* allele (*E1*) and also with one particular *het-d* allele (*D1*), but not with other alleles (**Figure 1B**). In other words, the other *het-d/e* alleles do not recognize *C2* as a ligand. To date, three alleles of *het-e* (*E1*, *E2*, and *E3*) are known to recognize *het-c*, while null variants are collectively known as the *e4* allele. Likewise, *het-d* has two known reactive alleles (*D1* and *D2*) and a non-reactive allele (*d3*). (In the literature, the different *het* alleles might be referred to as *het-E1*, *het-E2*, etc., but here we use a simplified terminology for readability).

Although the genetics of *het-d* and *het-e* are well understood, the precise characteristics of their sensor domain remain unclear, likely due to their repetitive nature and HIC properties. The original study that identified *het-e* sequenced the allele of the French strain A (here referred to as *E1*^A^) and reported 10 WD40 repeats (Saupe et al. 1995a). Later, Espagne et al. (2002) resequenced the same *E1*^A^ allele but found differences in multiple amino acids. They also sequenced the *het-e* allele of the strain C (*E2*^C^, 10 repeats) and used PCR and Southern blotting to estimate the number of repeats from several wildtype and mutant strains from the original French collection. On occasion, these two methods returned conflicting results, in which case they gave priority to the Southern blot analysis since PCR is susceptible to amplification artifacts (**Figure 1C**). Overall they concluded that at least 10 repeats are necessary for *het-e* to be reactive, that losing even a single repeat can break an allele, and that some alleles have the right size but are still non-reactive (*e4*) (Espagne et al. 2002; Saupe et al. 1995a). More recently, Chevanne et al. (2010) resequenced *E1*^A^ yet again and found it to contain 11 repeats instead. In the case of *het-d*, only a single allele has been sequenced, *D2*^Y^ (from the French strain Y), consisting of 11 full repeats and the first 30 amino acids of a 12th repeat at the C-terminus (Espagne et al. 2002). As for *het-e*, Espagne et al. (2002) examined the sizes of French wildtype *het-d* alleles by PCR and Southern blot, inferring that the functional *D1*^F^ also has 11 full repeats and a truncated one, while non-reactive alleles (*d3*) have either less or more repeats. Thus, the actual sequences of most active alleles remain unknown, precluding additional functional and evolutionary studies. Moreover, the reported sizes of functional HNWD alleles (e.g., 10 or 11) are at odds with the number of repeats expected from usual β-propellers, which is six to eight repeats but most often seven (Hu et al. 2017).

The growing availability of genomic data has thrust the study of fungal NLRs into new frontiers (Daskalov et al. 2015; Dyrka et al. 2014; Daskalov et al. 2020). However, modern whole genome sequencing using short-read (Illumina) technologies is not necessarily the solution for NLRs with HIC: tandem repeats with high sequence similarity can be notoriously difficult to assemble (Tørresen et al. 2019). The length of a single WD40 repeat is 126 bp (42 amino acids), close to the size of a typical Illumina read, making it a borderline case. Long-read technologies, such as PacBio or Oxford Nanopore Technologies (ONT), hold the promise of accurate genome assembly, especially as error rates and costs decrease (Sereika et al. 2022; Espinosa et al. 2024). Here we took advantage of published Illumina, PacBio, and ONT datasets of wildtype *P. anserina* strains (Vogan et al. 2019, 2021) to examine the HNWD alleles in the context of different sequencing technologies and assembly software. To resolve inconsistencies in the literature, we produced new ONT data to recover the reactive *het-d* and *het-e* alleles of the original French strains. Having established a reliable set of HNWD sequences, we assessed the interactions of a *het-e* allele seen in several wildtype strains. Finally, we discuss possible arrangements of their β-propeller domain using AlphaFold 3 protein structure models. Overall, we provide a basis for the study of the binding specificity and evolution of these variable immune receptors.

## Results

### The haplotypes of HNWD genes can be recovered consistently from long-read data but not from short-read assemblies

To assess the consistency of HNWD genes across sequencing efforts we first focused on three *P. anserina* strains: the Dutch strain Wa63+ and the French strains Y+ and Z+ (the + and -annotation signify the mating type). These three strains are haploid and their genomes were originally sequenced as paired-end (125 bp x 2, insert size ∼350) libraries with Illumina HiSeq 2500 at high coverage (>70x) (Vogan et al. 2019). In the same study, high-molecular-weight (HMW) DNA was also extracted from Wa63+ and Y+, which was then sequenced using either PacBio RSII or an R9 ONT flowcell, respectively (Vogan et al. 2019). Here, we re-sequenced these same strains using R10 ONT flowcells in a barcoded library (see Methods). Hence, this dataset allowed us to compare the HNWD haplotypes obtained from different sequencing technologies and assemblies of the same strains at different time points (**Table 1**).

**Table 1.**
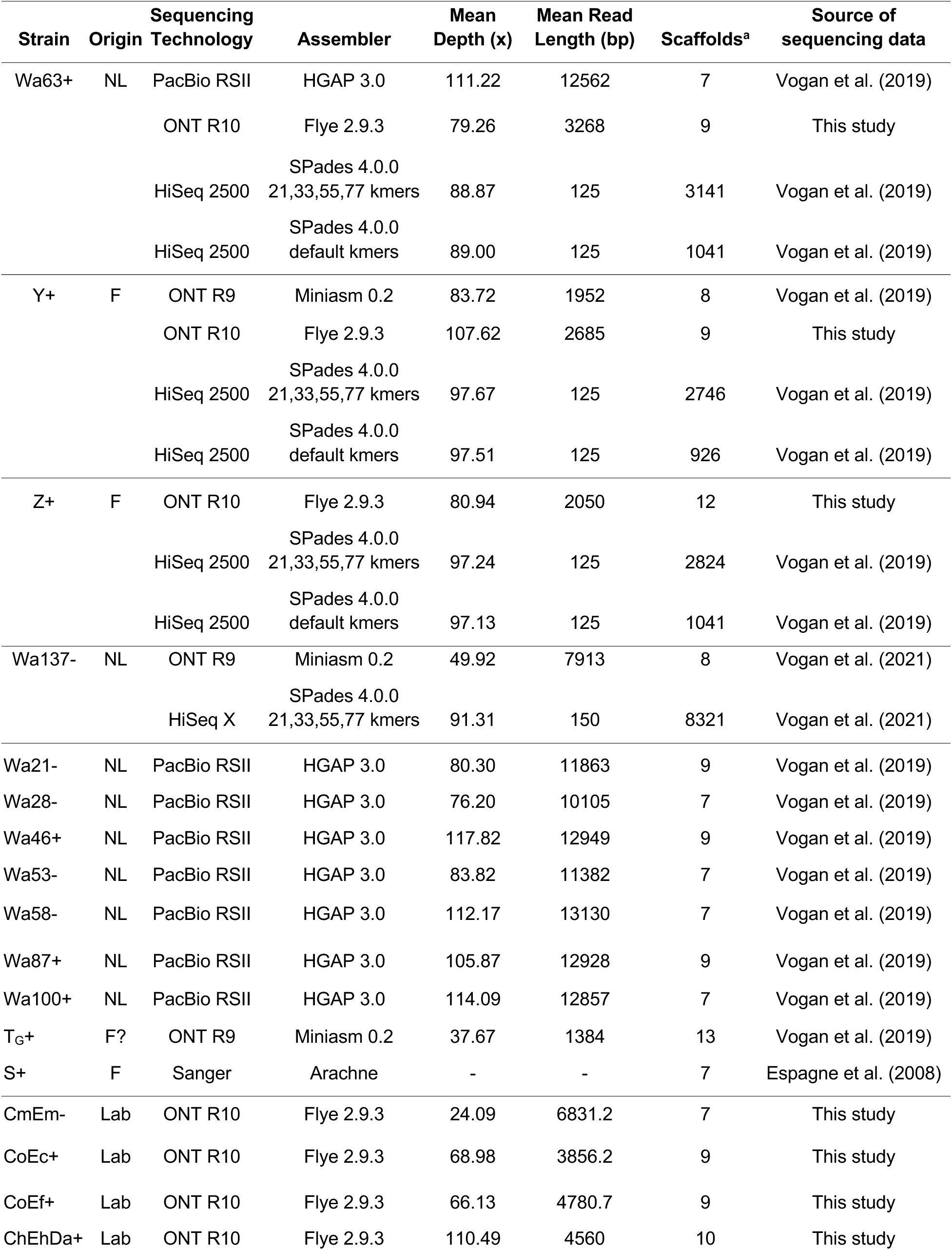
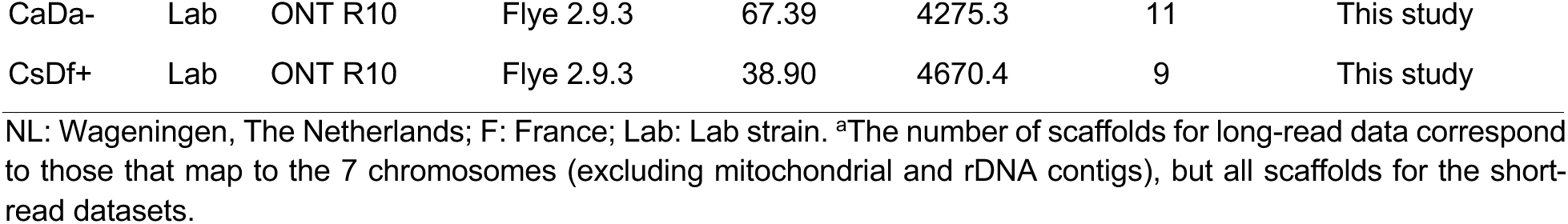
Whole genome assemblies of *P. anserina* strains used in this study.

We extracted the sequence of *het-d*, *het-e,* and *het-r* from each assembly and compared their WD40 domain (**Figure 2**). To facilitate the visualization, we classified the HIC repeats of each gene by assigning an arbitrary number based on unique combinations of seven amino acids at positions previously inferred to be under diversifying selection (Paoletti et al. 2007) (**Table S1**). In addition, we calculated dissimilarity scores among the HIC repeat classes based on an amino acid physicochemical dissimilarity matrix (Urbina et al. 2006). The scores were used to generate palettes in the CIE L*a*b* color space, such that similar colors imply similar physicochemical characteristics. We found that repeats were more different between paralogs than among the repeats of each paralog, so we assigned an independent palette per gene to facilitate contrast. See also **Figures S1**, **S2**, and **S3**.

**Figure 2.**
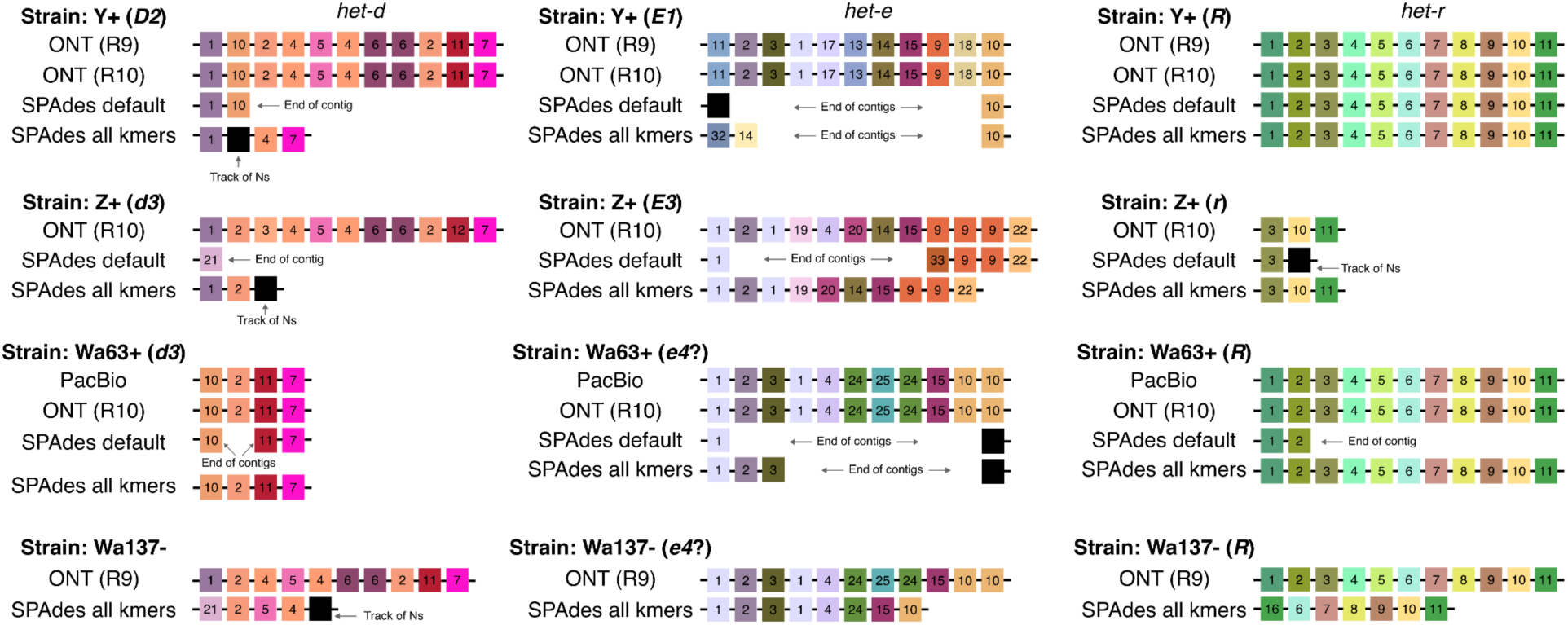
Assembly of the WD40 domain from different sequencing technologies. Only repeats with high internal conservation are shown. Each repeat was arbitrarily classified based on unique amino acid combinations, but the colors reflect their physicochemical similarity (each gene has an independent palette) Repeats with a track of missing data (Ns) are colored black. Black lines linking the repeats symbolize the containing scaffold.

We found that the long-read assemblies were in agreement, regardless of the technology and assembler. In the case of *het-r*, both Y+ and Wa63+ recovered the exact same sequence of repeats as reported for the reference *R* allele (Chevanne et al. 2010). This suggests that 1) the sequence obtained from the long-reads (and the reference itself) is correct, and 2) the HNWD alleles did not mutate despite an unknown amount of vegetative growth and culture transfers that the strains have undergone in the lab since isolation.

Most sequencing datasets of non-model fungal species are based on short-read data, implying that NLRs with HIC are usually assembled from Illumina reads. What is the likelihood that these assemblies are correct? As a proof of concept, we used the popular short-read assembler SPAdes (Prjibelski et al. 2020) to test if we can obtain equivalent haplotypes from the published Illumina data of Wa63+, Y+, and Z+. SPAdes constructs assembly graphs using multiple k-mer sizes, which can be selected automatically by the program (Prjibelski et al. 2020). In our case, the selected k-mer sizes were 21, 33, and 55 bp (“default” treatment). To promote contiguity in the assembly, we also produced assemblies using an additional larger k-mer of 77 bp (the “all kmers” treatment). As before, we extracted the WD40 domain of the HNWD genes but found that it is often fragmented into two scaffolds (**Figure 2**). In cases where an HNWD gene was fully contained within a scaffold, the haplotypes harbored tracks of missing data (Ns) or presented a different sequence than their long-read data counterparts (e.g., *het-e* in the strain Z+). The only exceptions where long (>5 repeats) alleles were correctly recovered from Illumina assemblies were those of *het-r* in the “all k-mers” SPAdes assemblies (**Figure 2**). Notably, some SPAdes assemblies included chimeric repeats that were not present in the wildtype long-read haplotypes (e.g., repeat variant 32 in the “all kmers” *het-e* sequence of the strain Y+).

The Illumina reads of Wa63+, Y+, and Z+ are relatively short, of 125 bp. Current Illumina technologies usually have slightly longer reads of 150 bp. To assess if that difference was enough to recover the HNWD alleles, we used published data of the strain Wa137-(Vogan et al. 2021). The genome of this strain was sequenced with R9 ONT flowcells as for Y+, but its short-read library was sequenced with the HiSeq X machine (150 bp paired-end, insert size ∼ 250 bp) (**Table 1**). For this read length, SPAdes defaults to all k-mers (21, 33, 55, and 77); hence we only evaluated the assembly with those parameters. As with the other strains, we found that the HNWD haplotypes recovered are shorter than the long-read assembly, omitting or creating repeats (**Figure 2**). We conclude from these analyses that HIC repeats are not recovered confidently from Illumina assemblies.

### Different WD40 repeat combinations result in the same functional allele

The molecular biology studies that first described *het-d* and *het-e* used lab strains constructed by backcrossing the alleles of French strains with known phenotypes into the genomic background of a reference strain (s, also referred to as “little s”) (Espagne et al. 1997, 2002; Chevanne et al. 2010; Saupe et al. 1995a). We sequenced the genome of some of these backcrossed strains using ONT R10 as above. The backcrossed strains are designated by their reactive genotypes. For example, the strain CmEm-contains the *het-c* (*C2*^M^) and *het-e* (*E2*^M^) alleles of the French strain M, while having the non-reactive *het-d* allele (*d3*^s^) of strain s. Likewise, the strain ChEhDa+ has the *het-c* (*C3*^H^) and *het-e* (*E1*^H^) alleles of the H strain and the *het-d* allele (*D1*^A^) of the A strain. The exceptions are strains with a null *het-c* allele, here termed Co (CoEc+ and CoEf+). The genome assemblies of these backcrossed strains consist of mostly full chromosomes or chromosome arms (**Table S1**).

The *het-e* and *het-d* sequences from these new genomes add to the collection of reliable sequences for alleles of known reactivity. The ChEhDa+ and CaDa-strains have the same *het-d* allele as the A strain (*D1*^A^), and the recovered sequences were identical, reinforcing the notion that long-read assemblies represent the real DNA sequence. The strain Z+ above belongs to the original collection of French strains with known phenotypes (Bernet 1967). The strain S (“big S”) is also part of this collection and its genome is considered the reference for the species, although it predates long-read technologies (Espagne et al. 2008). However, S has non-reactive *het-d* and *het-e* alleles that are relatively short (e.g., **Figure 1C**), and hence more likely to be correctly assembled. In addition, the strain Y+ is known to harbor a *D2* allele (Espagne et al. 2002) and an *E1* allele (L. Belcour, personal communication). Hence, current data allows preliminary comparisons of intra-allele variation (**Figure 3**).

**Figure 3.**
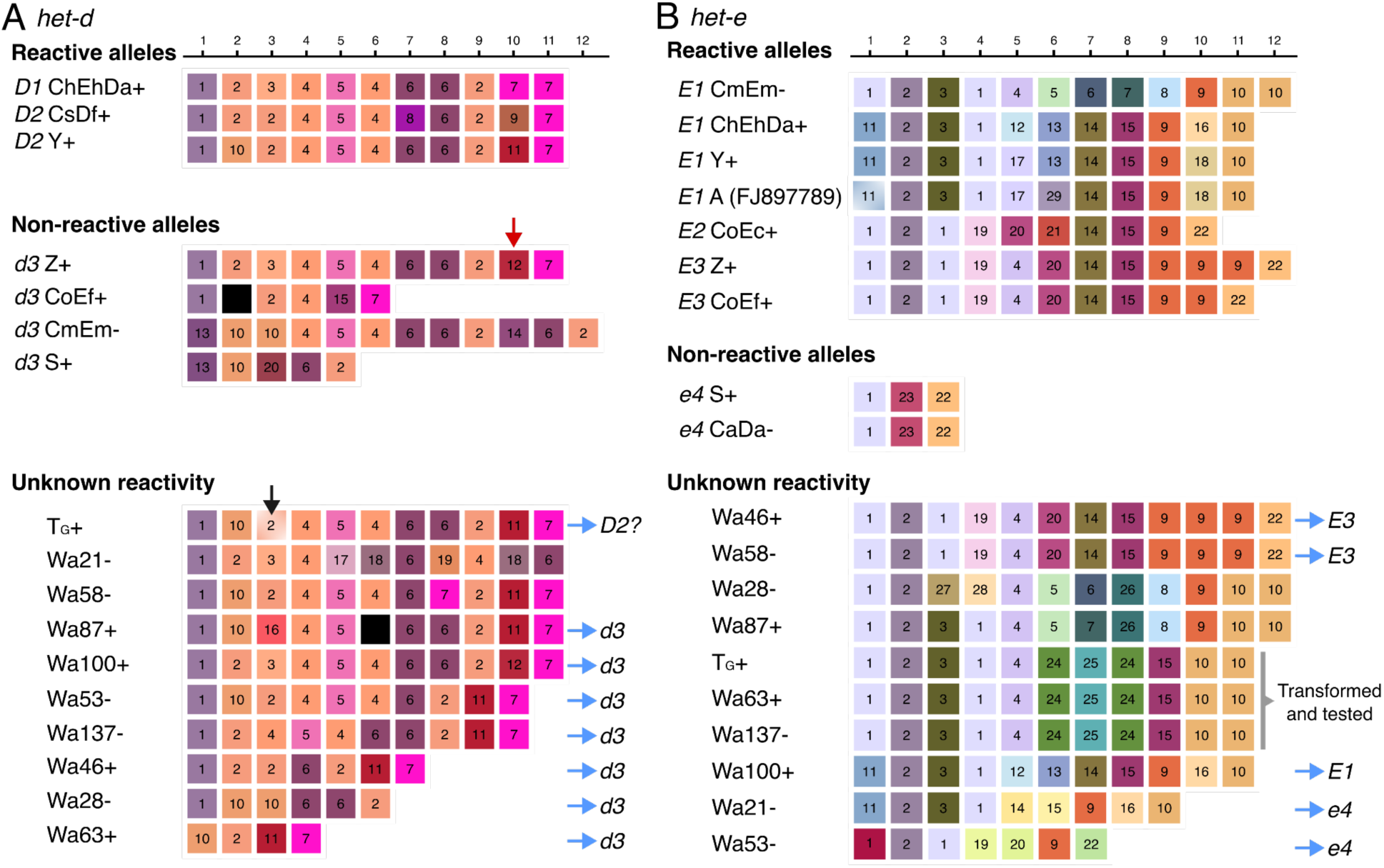
Long-read assemblies of *het-d* (**A**) and *het-e* (**B**) WD40 domain from different wildtype strains. Only repeats with high internal conservation are shown, arbitrarily classified based on unique amino acid combinations but colored based on their physicochemical similarity (each gene has an independent palette). Repeats containing stop codons or frameshifts are colored black. The red arrow highlights the single repeat distinguishing the reactive *D1* ChEhDa+ allele from the non-reactive *d3* Z+ allele. The black arrow marks the repeat with a deletion in the T_G_+ sequence that is likely a misassembly. Blue arrows point to inferred alleles based on sequence or the number of repeats. A specific allele of *het-e* was selected for phenotypic testing. The beginning of the first repeat is missing in the *E1*^A^ sequence (GenBank accession number FJ897789) but the missing amino acids happen to be perfectly conserved in all sequences and hence we inferred it to be identical to the first repeat of *E1*^Y^.

In agreement with previous reports, our long-read assemblies show that all the reactive *het-d* alleles have 11 repeats with HIC, while the reactive *het-e* alleles can be 10, 11, or 12 repeats long (**Figure 3**). However, the precise order and identity of the repeats from the published *D2*^Y^ and *E2*^C^ alleles show strong differences with the ONT assemblies (**Figure S4**). Likely, older methodologies had difficulties establishing the specific order of repeats, but the general inference that the *het-d* and *het-e* alleles with less than 10 repeats are non-reactive still holds (Saupe et al. 1995a; Espagne et al. 2002). Indeed, many of the non-reactive sequenced alleles are short (**Figure 3**). Nonetheless, as pointed out previously (Espagne et al. 2002; Saupe et al. 1995a), having the right number of repeats is not enough to create a reactive allele. A clear example is given by the non-reactive *het-d* allele of strain Z (*d3*^Z^), which is identical to the *D1*^A^ allele except for a single repeat at the 10th position (red arrow in **Figure 3A**). This repeat differs from its functional counterpart by just four amino acids (**Table S2**), and has a single amino acid difference with the repeat variant 11 at the same position in *D2*^Y^. The fact that some positions are highly conserved across allele classes further suggests that these positions are key for functionality (e.g., the 4th to 6th positions of *het-d* and second position of *het-e*).

Interestingly, none of the sequences from the same allele class are identical (e.g., all the different *E1* alleles), implying that although a single misplaced repeat can break an allele, there must be some flexibility at some repeat positions. For example, there is considerable intra-and inter-allele variation at the 5th and 6th repeat positions of *het-e* (**Figure 3B**). The *E1*^M^ allele (CmEm-) is particularly puzzling since it is quite different from the other *E1* alleles at the fifth to ninth repeats. Nonetheless, some positions might be diagnostic of an allele class. For example, all the *E1* alleles have the same repeat at the 3rd and 4th positions, as well as the same last repeat (regardless of the haplotype length), relative to the *E2* and *E3* alleles. Notably, the length polymorphism seems to be concentrated towards the last three repeats of the *E* alleles, with the *E3* alleles being the best illustration. In this case, a single repeat (classified as variant 9 in **Figure 3B**) is repeated in *E3*^Z^ relative to *E3*^F^. Likewise, the *E1*^M^ allele has an extra variant 10 repeat compared to the other *E1* alleles.

Having established a reference panel of allele sequences, we looked at published long-read data of other wildtype strains (Vogan et al. 2019, 2021). While T_G_+ and Wa137-were sequenced with ONT R9, the other strains were sequenced using the PacBio RSII technology (Vogan et al. 2019, 2021). From these strains, Wa100+ has the same *het-d* sequence as Z+, and hence has a *d3* allele (**Figure 3A**). The *het-d* sequence of T_G_+ is in fact very similar to that of *D2*^Y^, with the notorious exception of a single base-pair deletion at the third repeat (**Figure 3A**) and a substitution in position 17 (not under diversifying selection) of two repeats. Inspection of the short-read mapping to the assembly of this strain suggests that the deletion is a misassembly within a small homopolymer track (**Figure S5**), which is a more acute problem in R9 data than R10 or PacBio (Sereika et al. 2022). On the other hand, the strain Wa87+ has two stop codons in its sixth repeat that are supported by read mapping (**Figure S6**). Hence, this strain likely has a disrupted protein and can be tentatively assigned to a *d3* allele-type. All strains with less or more than 11 repeats can also be considered *d3* (Espagne et al. 2002). In the case of *het-e*, three strains have sequences identical to those in the reference panel: both Wa46+ and Wa58-have an *E3* allele, while Wa100+ has an *E1* allele (**Figure 3B**). Based on the similarity to *E1*^M^, the sequence of Wa87-might be *E1*, although that requires testing. Sequences with less than 10 repeats can be assigned to the *e4* allele.

### The *het-e* allele of Wa63+ does not recognize common *het-c* alleles

Although this is a small sample of strains, we noticed that one particular *het-e* sequence appeared in three wildtype strains: T_G_+, Wa63+, and Wa137-(**Figure 3B**). The origin of T_G_+ is unclear, although it might correspond to the French T strain (Vogan et al. 2019). The other two strains were both sampled in Wageningen, the Netherlands, but one in 1994 and the other in 2016. Hence, we wondered if this was an unidentified functional *E* allele. As these strains have *C2*, *C9*, and *C2* alleles, respectively, then these *het-e* sequences could not correspond to *E1* or *E2* alleles, as that would create self-incompatibility. To assess its reactivity, we cloned the *het-e* allele of Wa63+ on a plasmid and introduced it by transformation into two different recipient strains with no reactive *het-e* or *het-d* alleles: one with a *C1d3e4* genotype and another with a *C2d3d4* genotype (**Figure 4**). In this way, it is possible to assay incompatibility to the common *het-c* alleles (*C1*, *C2*, *C3,* and *C4*). In total, 24 transformants were tested in barrage assays against testers carrying the common *het-c* alleles. We found that all transformants were compatible with all *het-c* testers. In a control experiment, using a cloned *D1* allele introduced into a the *C1d3e4* recipient, 15 out of 24 tested transformants produced a barrage reaction. We conclude from this experiment that the *het-e* allele from Wa63+ does not lead to incompatibility with the common *het-c* alleles. Either this allele is inactive in incompatibility or, alternatively, it could lead to incompatibility to rare *het-c* alleles that were not tested in this experiment (see Discussion). Notably, this allele lacks a repeat variant 9, which is present towards the end of all known functional alleles.

**Figure 4.**
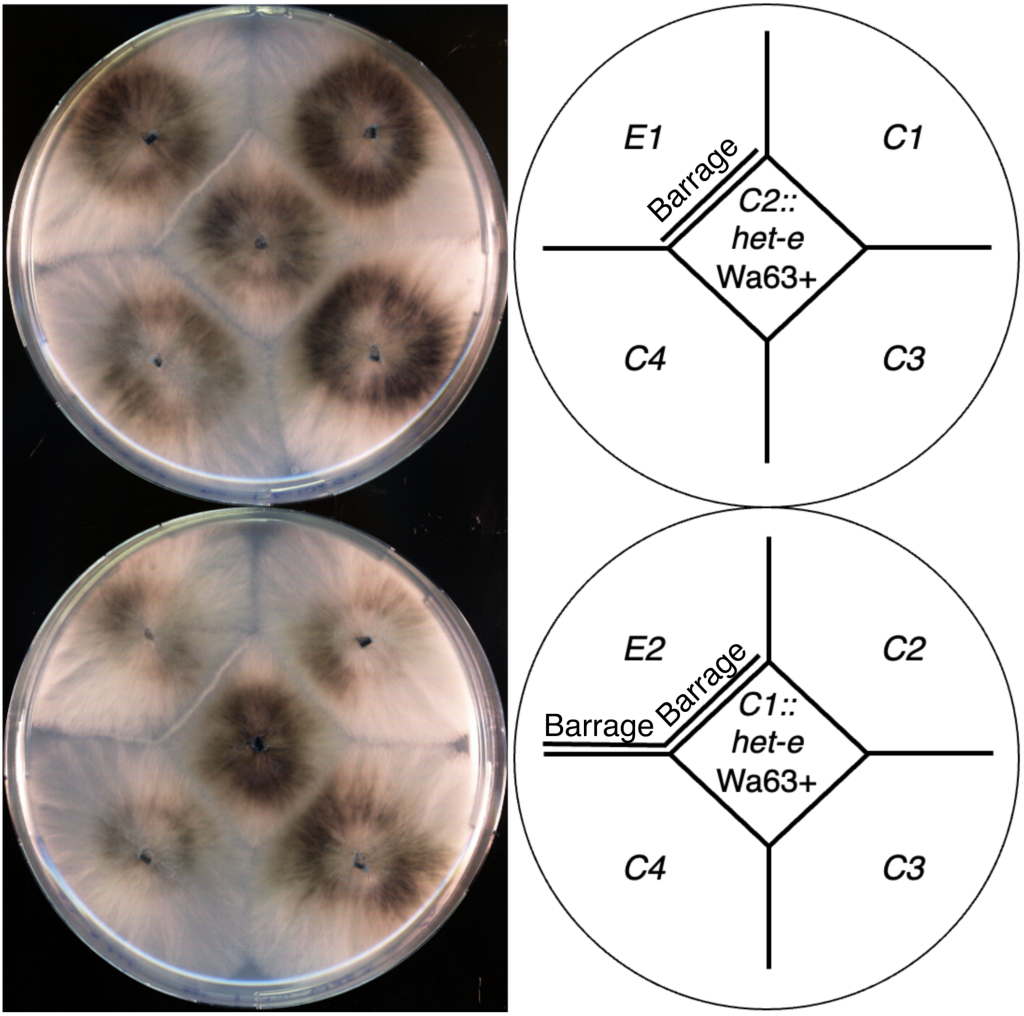
The *het-e* allele from Wa63+ is compatible with the four common *het-c* alleles. Barrage assay of a *C1* and *C2* strains transformed with the *het-e* allele of Wa63+ cloned on plasmid and tested with the four common *het-c* alleles. The *C2* recipient strain (upper plate) allows for testing against *C1*, *C3*, and *C4*. The *C1* recipient (bottom plate) allows for testing against *C2*, *C3* and *C4*. The *E1* and *E2* alleles (on the upper left on the upper and bottom plates respectively) are used for positive controls for the barrage (incompatibility) reaction. Note the barrage formation between the *E2* and *C4* testers in the bottom plate. In the strain designation, the *het-c*, *het-d* and *het-e* genotypes are omitted for clarity when strains carry inactive alleles.

### The reactive HNWD alleles likely form a double β-propeller structure with cryptic repeats

The WD40 β-propeller fold is formed by six to eight, but usually seven, units called “blades”, which arrange radially around a central tunnel (Fülöp and Jones 1999). Each blade, in turn, is formed by four antiparallel β-sheets named *a*, *b*, *c*, and *d*. By convention, a WD40 repeat does not exactly correspond to a blade, but instead starts with a *d* β-sheet from the previous blade, followed by *a*, *b*, and *c* sheets of the focal blade (**Figure S7A**). To close the circle, the last blade is often constructed from one to three β-sheets of the last repeat (the C-terminus), complemented by remaining β-sheets from the N-terminus, a configuration known as the molecular “velcro” (Neer and Smith 1996; Fülöp and Jones 1999).

Our results confirm that the most common reactive HNWD alleles display 11 HIC repeats, which would represent an atypical blade number. We turned to AlphaFold 3 to model the WD40 β-propellers of *het-e*. In a first experiment, we input a single repeat (the second HIC repeat in the *E2*^C^ allele) and modeled different combinations of blade numbers, from 6 to 9-mers. The AlphaFold model quality and confidence scores (pTM and ipTM, where 1 represents the best prediction) were highest for 7-mers and 8-mers (**Figure S7A**). As an alternative approach, we created an artificial sequence of 6, 7, 8, and 9 tandem identical repeats (**Figure S7B**). In this case, the 7-repeat sequence yielded the highest pTM value (0.95), suggesting seven is a relevant size for the *het-e* β-propellers.

Next, we modeled the HET-E1 WD40 domain on its own (**Figure 5**) and within the HET-C2/HET-E1^A^ protein complex (**Figure S8**). Although with relatively weak support scores (for WD40 domain only ipTM = 0.32 and pTM = 0.41; for full length ipTM=0.47 and pTM 0.59), the resulting multimer model is consistent with previous observations. The WD40 domain of HET-E1 is modeled as two independent 7-mer β-propellers (**Figure 5**), with the HET-C2 protein located at the predicted interaction surface that contains the hypervariable sites under diversifying selection (Paoletti et al. 2007) (**Figure S8**). The known specificity-defining residues of HET-C are also located in the predicted interaction surface (Bastiaans et al. 2014). Importantly, the second predicted propeller is made of four canonical highly conserved blades (8th to 11th) and three additional cryptic (divergent) blades located at the C-terminus of the protein, closed by a molecular velcro with a cryptic *d* β-sheet on the N-terminus (**Figure 5** and **Figure S8**). These modeling approaches suggest that active WD40 repeat domains have a mosaic structure with two propellers forming a clamp-like structure, one of which comprises a combination of four canonical and three cryptic repeats. Such a model provides a plausible explanation for the occurrence of an otherwise unusual number of repeats in active *het-d/e* alleles.

**Figure 5.**
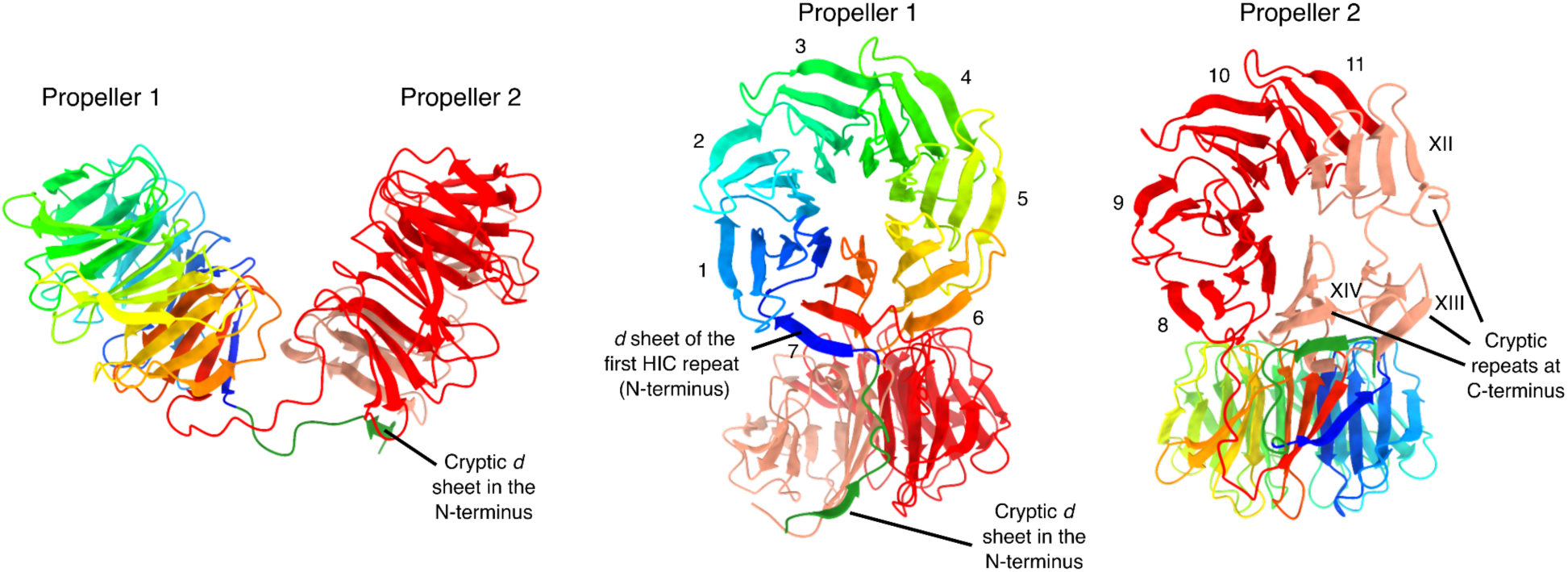
Ribbon diagrams of the WD40-domain structure from the HET-E1 protein (*E1*^H^ allele) produced by AlphaFold 3. The first 817 sites containing the HET and NACHT domains were removed for clarity. The first propeller is colored with a rainbow palette to illustrate the direction of the individual β-sheets. The cryptic *d* β-sheet in the N-terminus of the WD40 domain that forms the molecular velcro with the C-terminus is also highlighted (forest green). The second propeller is colored based on HIC (red) and cryptic (salmon) repeats. Individual blades are numbered with Latin (HIC) or Roman (cryptic) numerals.

## Discussion

Evolutionary and functional research of immune system genes, including NLRs, often comes with technical and methodological challenges. For example, rapid evolution might limit phylogenetic reconstructions or ascertainment of homology (Messier-Solek et al. 2010). Likewise, the presence of paralogs and association with transposable elements can result in fragmented genome assemblies at precisely the NLR locations (Yuen et al. 2014; Tørresen et al. 2019). Thus, highly similar repeats at the C-terminal domain (i.e., HIC) can act as the final nail in the coffin for the assembly of certain NLRs. Here we show that current long-read technologies fully overcome that problem for the WD40 domain of HNWD genes, regardless of the software and the technology used. By sequencing the alleles used in classical genetic studies, we provide confidence to older inferences on the characteristics of reactive alleles, but highlight the potential inconsistencies of Illumina-derived assemblies in general. These high-confidence sequences, in turn, can be used to infer the phenotype of other strains, bypassing difficult and time-consuming lab experiments. Ultimately, having multiple alleles that display the same phenotype allowed us to showcase both the fragility and the flexibility of β-propellers as sensor domains, properties that might be advantageous for immune receptors.

Classic experiments designed to inactivate *het* genes found that the HNWDs have much higher mutation rates than their binding partners, *het-c* and *het-v* (Labarère 1973). Subsequent studies demonstrated that the HNWD genes are particularly susceptible to mutations at their sensor domain, altering or inactivating their recognition specificities by losses, gains, and shuffling of repeats (Chevanne et al. 2010; Espagne et al. 2002; Bastiaans et al. 2014). This “repeat instability” led to the suspicion that HNWDs (and potentially all NLRs with HIC) might easily break during somatic growth, driven by unequal crossing-overs and interparalog recombinations (Chevanne et al. 2010; Paoletti et al. 2007). Our results suggest that if this process occurs during vegetative propagation in the lab, it is too infrequent in the sequencing reads to influence the assembly graph. Detecting somatic repeat mutations may require extremely deep long-read sequencing efforts. Nonetheless, intra-thallus diversity in NLR repeats might still be subject to selection in nature, in particular if occasional variants happen to improve recognition of nonself. It has been suggested that fungal NLRs might have a general innate immune system function similar to that of plants and animals (Uehling et al. 2017; Paoletti and Saupe 2009). Immune system genes often display high diversity maintained by balancing selection, which is also a characteristic necessary for genetic systems controlling conspecific self-nonself recognition (allorecognition) (Aanen et al. 2008; Buckley and Dooley 2022). Heterokaryon incompatibility is a type of allorecognition, and NLRs might occasionally get co-opted as *het* genes in different fungal lineages (Paoletti and Saupe 2009). From that perspective, the high mutational rate associated with HIC repeats might be advantageous for both the innate immune system and allorecognition functions.

The fact that the Wa63+ *het-e* allele contains 11 HIC repeats, the usual size of a functional allele, and occurs in three different strains, sampled in different places and years, suggested that this might not be a random spontaneous mutant allele. However, our transformation essays demonstrated that this allele does not trigger an incompatibility reaction with the most common *het-c* alleles. That leaves us with three possibilities: 1) the allele is truly nonfunctional and its frequency is maintained by genetic drift; 2) the allele can only recognize some rare *het-c* alleles that were not tested here; or 3) the allele can recognize another ligand, such as a pathogen-derived molecule. Theoretically, an NLR *het* gene could simultaneously retain an ancestral immune function, further contributing to the maintenance of genetic diversity (Aanen et al. 2008; Bastiaans et al. 2014). Population and ecological studies on *Podospora* NLRs might help clarify this point.

Sequencing multiple versions of the same *het* alleles revealed a surprising diversity of repeat numbers and sequence combinations, implying that the mutational input can easily converge to the same phenotypes. The sequenced *E1* and *E3* alleles can either be 11 or 12 HIC-repeats long, while the one known *E2* allele has 10 HIC repeats. AlphaFold 3-predicted models were consistent with the idea that the WD40 domain of HNWD genes folds into two β-propellers and revealed the presence of cryptic repeats at the C-terminus. As several length differences between *het-e* variants of the same phenotype occur at the end of the HIC region, perhaps the second β-propeller can potentially be formed by 6 to 8 blades (in combination with the cryptic repeats) and remain functional. In contrast with the flexibility observed in *het-e*, the three known functional alleles of *het-d* all have 11 HIC repeats, and a single repeat change in the second propeller completely inactivated the *het-d* allele of the strain Z. Likewise, the only known reactive allele of *het-r* has 11 HIC repeats (Chevanne et al. 2010). One might speculate that the flexibility of *het-e* is related to the exact form of the cryptic repeats, which correspond to a region highly diverged between the HNWD paralogs.

The presence of highly similar repeats in the WD40 sensor domain of HNWD genes might be peculiar but is not a unique case for WD40 proteins. A large-scale screening of proteins with WD40 domains (not just NLRs) across the Tree of Life revealed that HIC happens most often in fungal and bacterial genomes (Hu et al. 2017). Moreover, NLRs with other types of superstructure-forming domains, such as ankyrin, tetratricopeptide, and HEAT repeats, are also known to have HIC in different fungal groups (Dyrka et al. 2014; Daskalov et al. 2020). There is even a report of a leucine-rich repeat NLR with HIC in a sea urchin genome (Hibino et al. 2006). In all these cases, the size of these repeat types is very similar to those of the WD40 repeats, between 24 and 42 amino acids (Yoshimura and Hirano 2016; Gupta and Chahota 2024; Marold et al. 2015). Therefore, the challenges we faced with short-read assembly of HNWD alleles are likely to apply to other NLRs across various taxonomic groups.

## Conclusion

Long-read technologies have been instrumental in the correct assembly of plant and animal NLRs since their development (e.g., Witek et al. 2016; Tørresen et al. 2018). However, the study of fungal NLRs is relatively new, and most genomic resources used in previous analyses have been Illumina-based (Daskalov et al. 2020), simply because most non-model species lack high-quality assemblies. Certainly, many aspects of NLRs can be fully studied from Illumina data, such as domain composition, diversity, and phylogenetic relationships. However, functional molecular biology studies can only do so much without high-confidence sequences, as in the case of *het-d* and *het-e*. Despite the availability of short-read population genomics data, the allele frequencies of these HNWD genes remain unknown. Such a gap hinders the study of potential balancing selection forces acting on them (Ament-Velásquez et al. 2022). The increased availability of long-read assemblies will not only address this limitation, but will simultaneously allow for the study of other aspects of their biology. For example, the genomic location of an NLR might influence its epigenetic modifications or mutation load (Sutherland et al. 2024). Looking forward, comparative studies of HNWD genes across populations and species, coupled with functional assays, may uncover novel roles for these genes beyond heterokaryon incompatibility. Additionally, integrating structural predictions with mutational analyses can clarify how β-propeller architecture contributes to the specificity of these immune receptors.

## Materials and Methods

### Fungal material and culture conditions

The strains used in this study were obtained from either the University of Bordeaux (Saupe et al. 1995a) or from the collection maintained in the Johannesson Lab at Stockholm University, which in turn came from the Laboratory of Genetics at Wageningen University (van der Gaag et al. 2000; Vogan et al. 2019). Work with all strains was done using monokaryotic (haploid) isolates, including those corresponding to the sequenced monokaryons in Vogan et al. (2019, 2021). Hence, strains are designated by their name and their mating type (e.g., Wa63+ is the Wageningen Collection strain 63 with a mating type +).

Mycelia for DNA extraction was obtained from two sources: Petri dishes (strains Y+, Wa63+, and Z+) and liquid cultures (all the strains with introgressed *het-c*, *het-d,* and *het-e* alleles into the strain s background). The cultures on Petri dishes were done with HPM media (Vogan et al. 2019) plates topped with cellophane disks cut from X50 Cellophane membrane 14x14 cm sheets (Fisher Scientific GTF AB, product code 11927535) and previously autoclaved in deionized water between filter paper disks (Cassago et al., 2002). Plates were incubated at 27°C under 70% humidity for a 12:12 light:dark cycle for two or three days (if left longer the mycelia ages and becomes harder to remove from the cellophane). Around 100 mg of mycelia were harvested by scraping the cellophane disk with a cell scraper (Sarstedt, Inc., 83.3951) and stored at -70°C.

It has been reported that *P. anserina* cultures in Luria-Bertani broth (LB) do not undergo senescence and are appropriate to get abundant and healthy mycelia (Benocci et al. 2018). We tested the use of both Luria-Bertani agar (LA) plates and LB cultures for mycelia harvesting. We found that the growth in LA or LB media significantly varies depending on the *P. anserina* strain. While strains S (used by Benocci et al.) and s thrive, some strains from the Wageningen collection exhibit poor growth. We also found that LA plates are not appropriate for the mid-term storage of strains. Hence, we used LB cultures just for the DNA extraction of the *het*-gene introgressed strains. Specifically, we used a modified LB recipe from Benocci et al. (2018) that contains 10 g/L Tryptone, 5 g/L Yeast extract, 5 g/L NaCl, and 0.02 g/L Thymine 99%, to which we added biotin and thiamin to a final concentration of 5 μg/L and 100 μg/L, respectively, and 1 mL/L of the trace element solution of van Diepeningen et al. (2008). We cut pieces of agar with mycelium from PASM0.2 plates (van Diepeningen et al. 2008) grown as for the HPM plates above, and used them as inocula for flasks containing 200 mL of modified LB. We incubated the flasks at 27°C and 120 RPM for five days (Vogan et al. 2019). The resulting mycelia balls were recovered from the flask with sterilized tweezers and stored at -70°C before extraction.

### DNA extraction and sequencing

Whole-genome DNA was extracted with the Zymo Quick-DNA Fungal/Bacterial Miniprep Kit D6005 (Zymo Research; https://zymoresearch.eu/) and quantified with a Qubit 2.0 Fluorometer (Invitrogen). For the strain CmEm-, ∼800mg of mycelia were used for high-molecular-weight DNA extraction using the QIAGEN Genomic-tip 100/G kit (Qiagen).

ONT sequencing was performed in-house using a Native Barcoding Kit 24 V14 SQK-NBD114.24 and a MinION Mk1C machine following the standard protocol. In total, 12 strains were barcoded into two pools (pool1: CmEm-, CoEc+, CoEc-, Y+, Z+, and Wa63+ with barcodes 1 to 6, and pool2: CoEf+, ChEhDa+, ChEhDa-, CaDa-, CsDf+, and CsDf-with barcodes 7 to 12). Each pool was sequenced in two separate R10.4.1 flow cells (FLO-MIN114), aiming at loading around 10-20 fmols of library for optimal duplex output (while assuming a highly fragmented DNA extraction to the detriment of sample CmEm-). Both libraries included 1 μl of diluted DNA control sample (DNA CS), a 3.6 kb standard amplicon used to QC the library. We added 5 μl of Bovine Serum Albumin (Invitrogen UltraPure BSA 50 mg/ml, AM2616) to the flow cell priming mix as recommended. The four flow cells (first two for pool1 and last two for pool2) were run until about 50 pores remained active (for 41 to 54 hours), generating 10.09 Gb,

9.89 Gb, 9.19 Gb, and 9.06 Gb estimated bases, respectively. All runs had the following settings: pore scan frequency of 1.5 hrs, minimum read length of 200 bps, read splitting on, and active channel selection on. The strains CoEc-, ChEhDa-, and CsDf-yield identical results to their opposite mating type counterparts so they were not discussed further in this study.

PCR Amplification of *het-e* in **Figure 1** was done with the forward 5’-GCCCTTGTATTTGCACCGAC-3’ and reverse 5’-CGTCCTGAGTAACAGCCAAGAAC-3’ primers, using the following temperature regime: 95 °C for 1 min; 35 cycles at 95°C for 15 s, 64°C for 15 s, and 72°C for 30 s; and 72°C for 7 min. The PCR reaction contained 8 μl ddH20, 0.5 μl of each primer (10 μM), 1μl of sample DNA, and 10 μl of MyTaq Red Mix (Meridian Bioscience™) for a total volume of 20 μl.

### Basecalling

During sequencing in the MinION Mk1C machine, we activated the “Fast model, 400 bps” for live basecalling with guppy v7.1.4 (embedded in MinKNOW v23.07.12). These reads were used only for preliminary coverage assessment per sample and automatic demultiplexing. The demultiplexed pod5 files were basecalled using Dorado v. 0.5.3 (https://github.com/nanoporetech/dorado/) with the dna_r10.4.1_e8.2_400bps_sup@v4.3.0 model. The resulting BAM files were transformed into fastq files with the bam2fq program of SAMtools v. 1.19.2 (Danecek et al. 2021). Reads corresponding to the DNA Control Sample (DNA CS) introduced during library preparation were removed using chopper v. 0.7.0 (De Coster and Rademakers 2023).

### Genome assembly and sequence analyses

For each sample, we removed reads that contained perfect matches to ONT native barcodes assigned to other samples (0.06% to 0.26% of the reads). We removed barcodes and performed minimum quality control with fastplong v. 0.2.2 (Chen 2023) and parameters –trimming_extension 20 -l 50 -q 15 -d 0.1 (hereafter, cleaned ONT reads). The cleaned ONT reads of each sample were used as input for Flye v. 2.9.3 (Kolmogorov et al. 2019), with parameters --nano-hq --iterations 2. The scaffolds were oriented to match the chromosomes of the reference genome Podan2 (Espagne et al. 2008). We visually looked for major rearrangements by mapping all assemblies to Podan2 with the NUCmer program of MUMmer v. 3.23 (Kurtz et al. 2004). The Integrative Genomics Viewer (IGV) browser was used for read-mapping visualization (Thorvaldsdóttir et al. 2013). Median read length and depth of coverage of the ONT R10 datasets were estimated by mapping the cleaned reads to their respective assemblies using minimap2 v. 2.26 (Li 2018) and feeding the produced BAM file to Cramino v. 0.14.1 (De Coster and Rademakers 2023). Equivalent values for published long-read assemblies of Wa63+ and Y+ genomes were taken from Vogan et al. (2019).

The paired-end Illumina reads of the strains Wa63+, Y+, Z+, and Wa137-were retrieved from NCBI’s Sequence Read Archive (accession numbers SRX5458088, SRX5458091, SRX11405146, and SRX8537866) and assembled with SPAdes v. 4.0.0 (Prjibelski et al. 2020) using the --careful parameter and either the default k-mers setting (Wa63-, Z+, and Y-) or the k-mers 21, 33, 55, and 77 (all strains). The Illumina reads were mapped back to each assembly using BWA v. 0.7.18 (Li and Durbin 2009). The resulting BAM file was sorted with SAMtools v. 1.21 (Danecek et al. 2011) and the duplicates were marked with Picard v. 3.3.0 (http://broadinstitute.github.io/picard/) with a value of 100 (Wa63-and Y-) or 2500 (Wa137-) for the --OPTICAL_DUPLICATE_PIXEL_DISTANCE parameter. Finally, the deduplicated BAM file was given as input of Qualimap v. 2.2.2d to obtain the average depth of coverage per assembly.

The nucleotide sequences of *het-d*, *het-e*, and *het-r* were extracted from each assembly using the script query2haplotype.py v. 2.22 available at https://github.com/SLAment/Genomics with parameters --haplo --extrabp 800 --minsize 400 --vicinity 15000 --identity 95 and the S+ allele as query. The sequences were manually aligned and given as input for a custom snakemake pipeline for WD40 repeat classification (https://github.com/SLAment/FixingHetDE). We employed a REGEX string to identify each repeat, defined as in Hu et al. (2017), and used the amino acids 10, 11, 12, 14, 30, 32, and 39 for classification based on their high dN/dS ratios (Paoletti et al. 2007).

Pairwise physicochemical dissimilarities between the different repeat variants were calculated based on the same seven high dN/dS positions by summing the pairwise distances at each position, as given by the amino acid physicochemical dissimilarity matrix in (Urbina et al. 2006). To generate color palettes for displaying the repeats, colors were chosen in the three–dimensional CIE L*a*b* color space (CIE 2019) using a variant of non-metric multidimensional scaling (NMDS) which matches the relative physicochemical distances of the repeat variants as closely as possible to the to the relative perceptual distances of the colors, while constraining the output to colors which can be represented in the sRGB gamut. This palette generation algorithm was inspired by Gecos (Kunzmann et al. 2020), and was implemented as a custom R script (available at https://github.com/SLAment/FixingHetDE) using the Python source for Gecos (https://github.com/biotite-dev/gecos) as reference.

### Cloning and transformation of *het-e*

The *het-e* Wa63+ gene was PCR-amplified with oligonucleotides osd192 (CAAGGTTGTGGCGGTTTCAG) and osd193 (GCGTTTGACAAGACGGTGAC) (respectively positioned at 605 nt upstream and 459 nt downstream of the ORF) on 33 ng of genomic DNA extracted from the Wa63+ strain using the Q5^®^ High-Fidelity 2X Master Mix (New England Biolabs, M0492S). We cloned 50 ng of PCR product in a pCR-Blunt II-TOPO^®^ vector using a Zero Blunt TOPO^®^ PCR Cloning Kit (Invitrogene, 45-0245) according to the manufacturer protocol. The ligation reaction was diluted at 1:4 in water and chemically NEB^®^ 5-alpha Competent *E. coli* (New England Biolabs, C2987H) were transformed with a twelfth of the ligation reaction.

DNA transformation was performed as previously described (Bergès and Barreau 1989) using the p1 vector, a pBlueScript-II derived vector containing the *nat1* nourseothricin acethyl transferase gene in co-transformation and using a 5 ug of the *het-e*-bearing plasmid and 1 μg of the p1 co-transformation vector. The recipient strains for transformation were *C2d3e4* (*het-c2 het-d3 het-e4*) and *C1d3d4* (*het-c1 het-d3 het-e4*). Five days after transformation, 24 individual transformants in each transformation were tested in barrage assays against the four common *het-c*-alleles on corn meal agar.

### Prediction of protein structure

We used the server of AlphaFold 3 (Abramson et al. 2024) available at https://alphafoldserver.com/ to model the protein structure of the WD40 domain. We used the protein sequence of the second HIC repeat in the *E2*^C^ allele (TGTQTLEGHGGSVWSVAFSPDGQRVASGSDDKTIKIWDAASG) as an arbitrary representative of a typical *het-e* HIC repeat. We gave this repeat to AlphaFold in 6, 7, 8, and 9 copies (multimers) to assess their predicted structure with default parameters. In addition, we input the protein sequence of HET-C2 (GenBank accession number AAA20542.1) and HET-E1^A^ (FJ897789; **Figure S5**) or HET-E1^H^ (**Figure 5**), also with default parameters. Only the first predicted model (number 0) was considered. We visualized protein structures in USCF ChimeraX v1.8 (Meng et al. 2023).

## Data availability

The snakemake pipeline used for WD40 repeat classification, as well as the all the nucleotide sequences of the het genes (aligned and in fasta format) are available at https://github.com/SLAment/FixingHetDE. All genome assemblies generated in this study have been submitted to the Dryad Digital Repository (https://doi.org/10.5061/dryad.h18931zww).

## Declaration of generative AI and AI-assisted technologies in the writing process

During the preparation of this work, the authors used ChatGPT to improve the flow and grammar of some parts of the manuscript. After using this tool, the authors reviewed and edited the content as needed and take full responsibility for the content of the published article.

## Acknowledgments

We thank María de la Paz Celorio Mancera, Anbar Khodabandeh, Jennifer Molinet, and Javier Pinto for their assistance and support in the laboratory, as well as Kalle Tunström for advice with ONT sequencing. This work was supported by the Swedish Research Council (grant 2022-00341) and the Stiftelsen Anna-Greta och Holger Crafoords fond (CR2023-0039) to S.L.A.-V. The computations were performed on resources provided by NAISS at Uppsala Multidisciplinary Center for Advanced Computational Science (UPPMAX) partially funded by the Swedish Research Council through grant agreements no. 2022-06725 and no. 2018-05973, under projects NAISS 2024/23-530 and NAISS 2023/22-924. Molecular graphics and analyses performed with UCSF ChimeraX, developed by the Resource for Biocomputing, Visualization, and Informatics at the University of California, San Francisco, with support from National Institutes of Health R01-GM129325 and the Office of Cyber Infrastructure and Computational Biology, National Institute of Allergy and Infectious Diseases.

## Supplementary Figures

**Figure S1.**
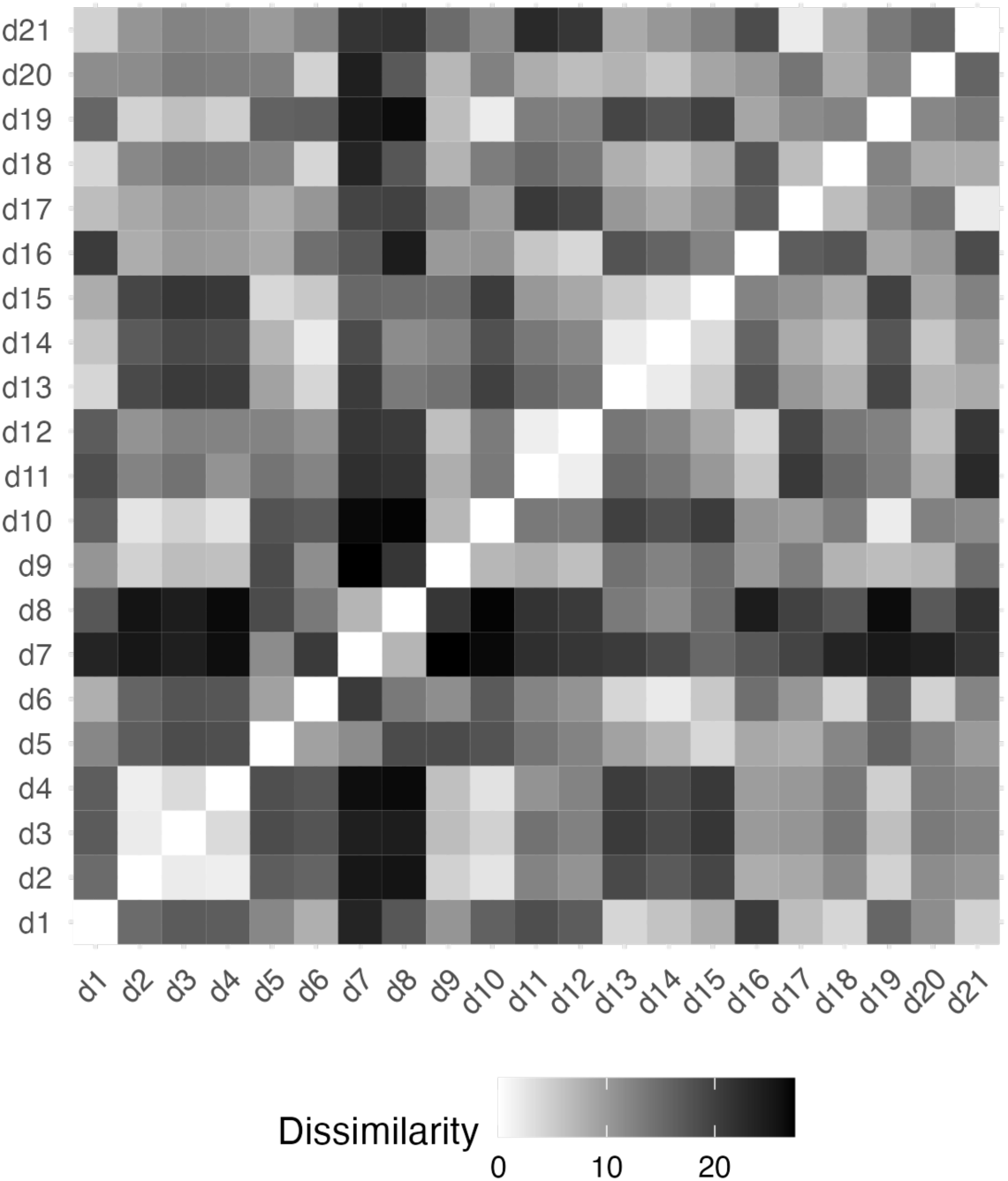
Heatmap of dissimilarity between the classes of HIC WD40 repeats of the *het-d* gene, based on seven amino acids.

**Figure S2.**
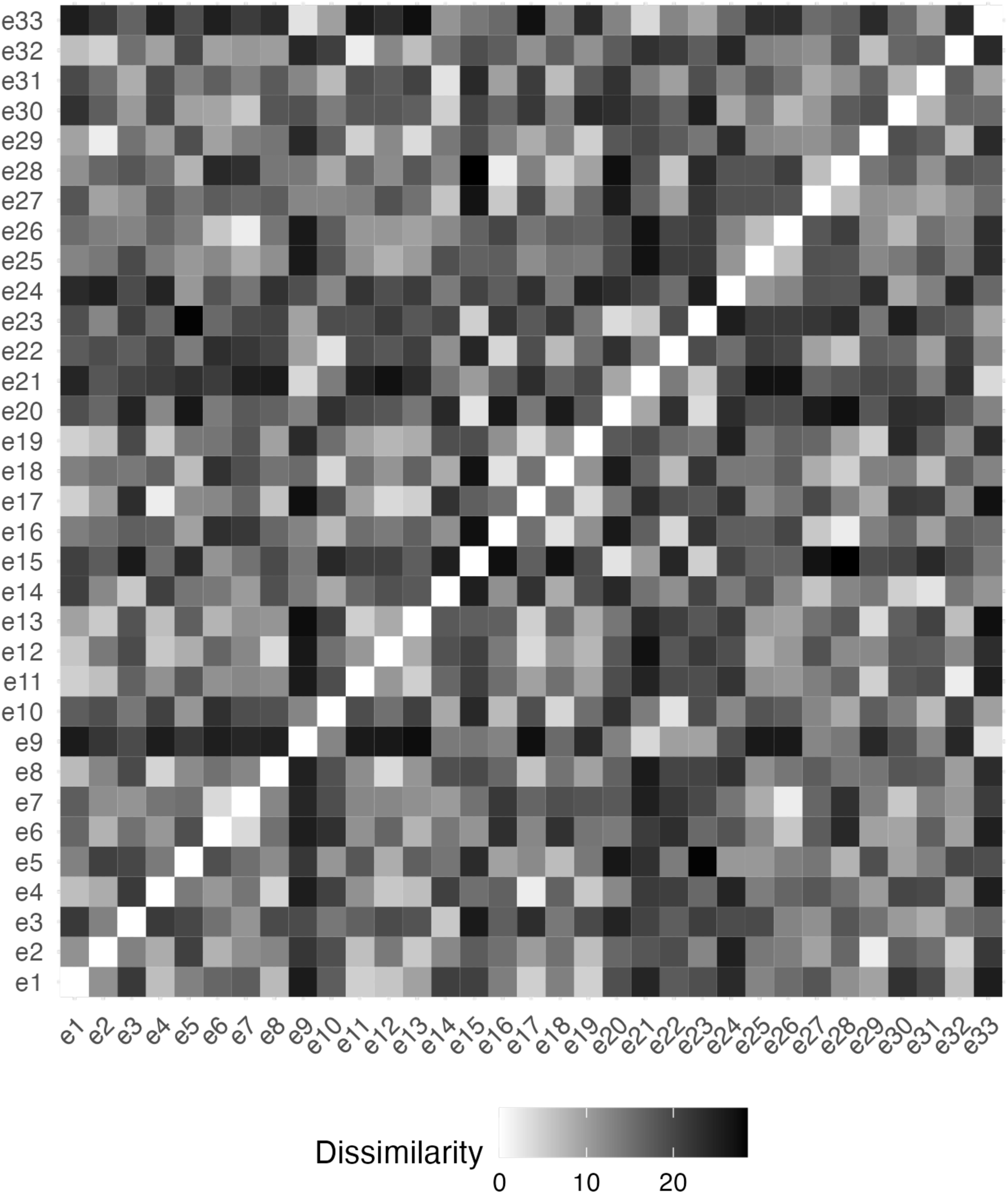
Heatmap of dissimilarity between the classes of HIC WD40 repeats of the *het-e* gene, based on seven amino acids.

**Figure S3.**
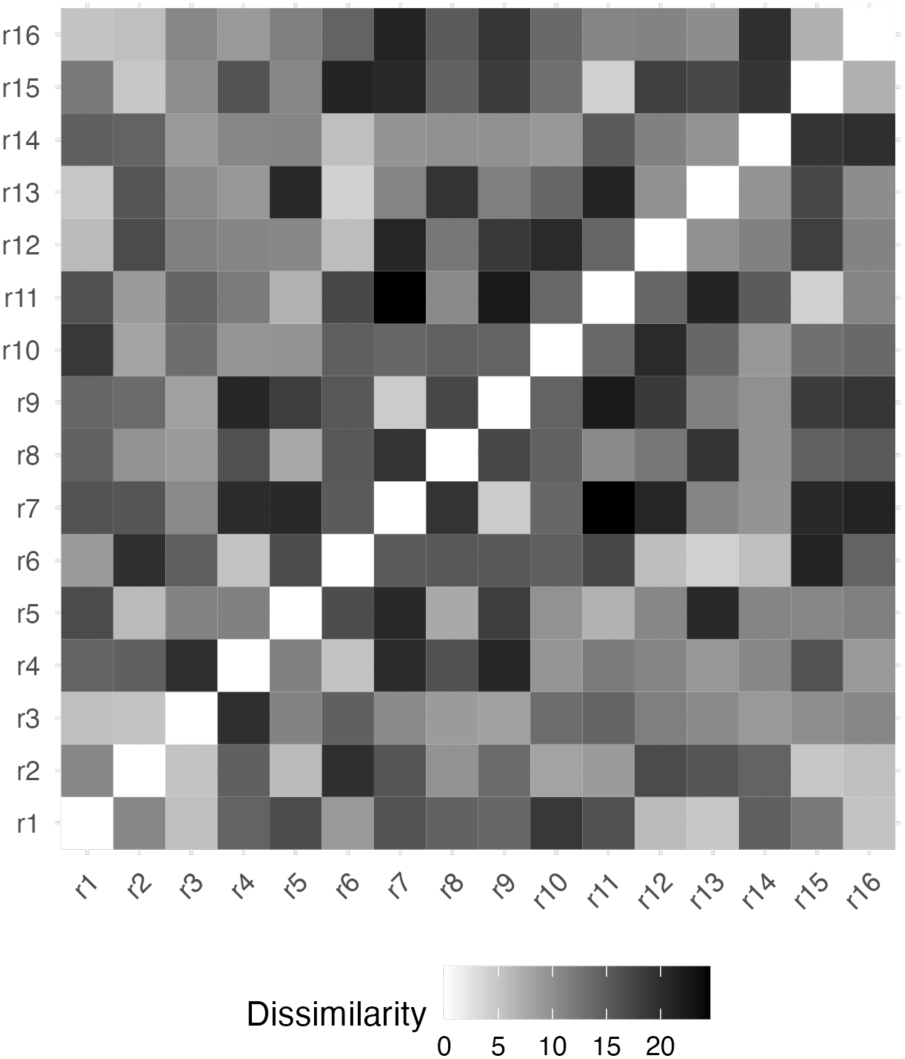
Heatmap of dissimilarity between the classes of HIC WD40 repeats of the *het-r* gene, based on seven amino acids.

**Figure S4.**
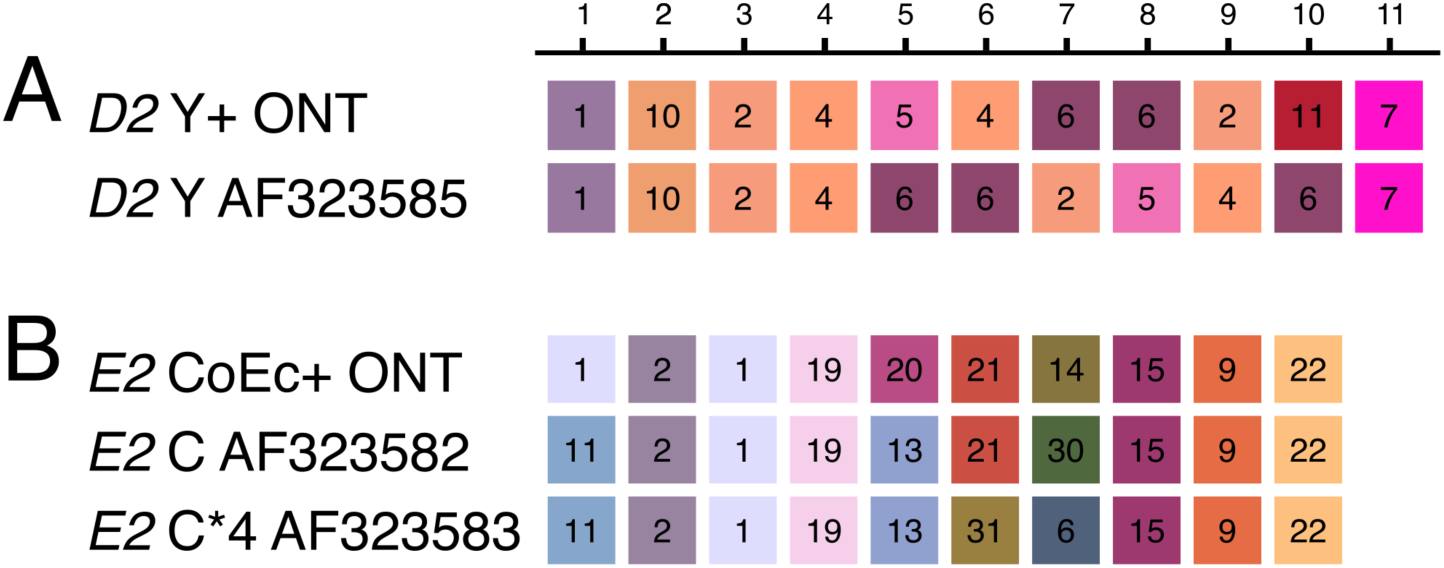
Comparison between the published alleles *D2*^Y^ (**A**) and *E2*^C^ (**B**) and corresponding long-read assemblies. The *E2* C*4 allele is a mutant of the original *E2*^C^ allele as reconstructed in the original study of Espagne et al. (2002).

**Figure S5.**
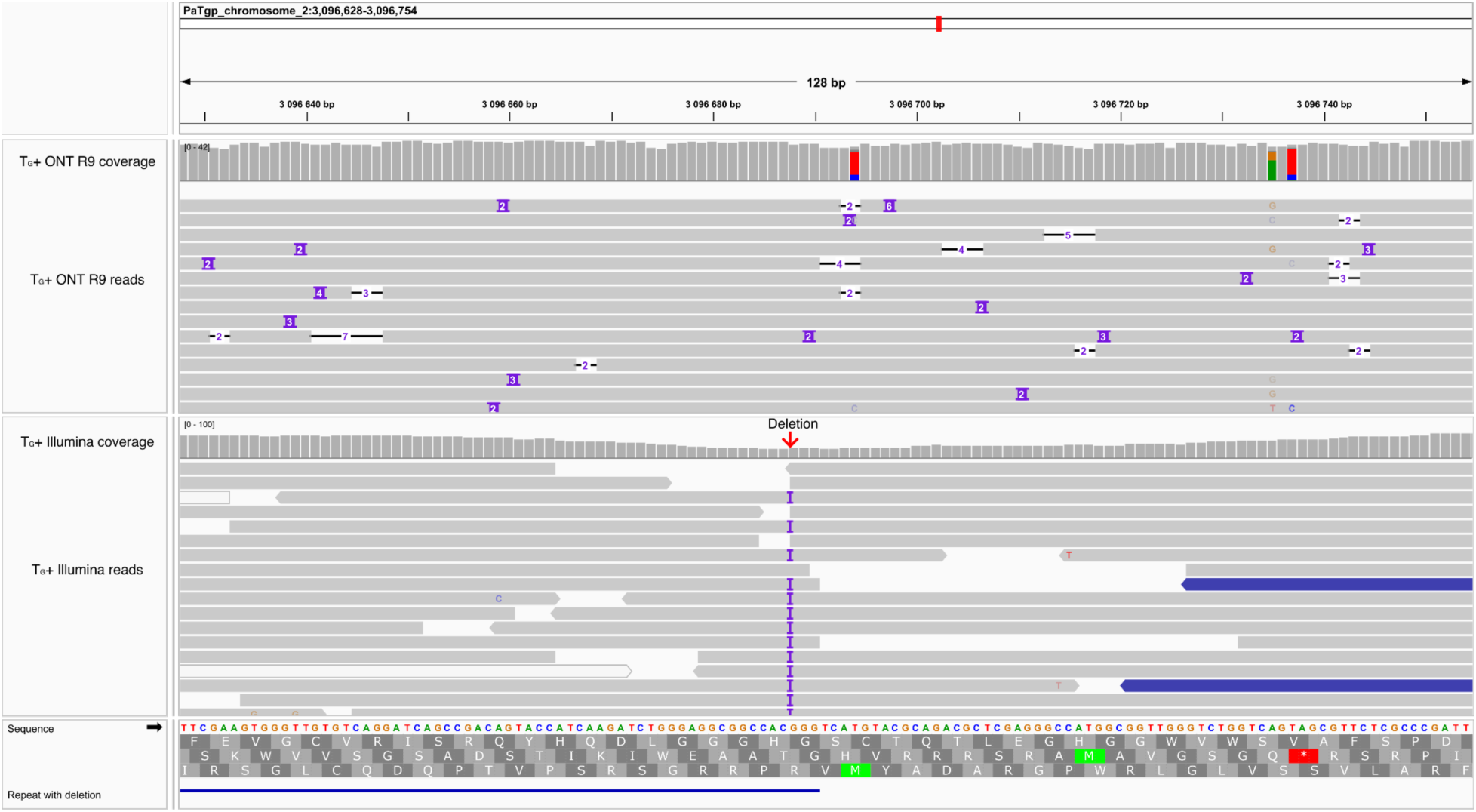
Short-and long-read mapping of the strain TG+ displayed in the Integrative Genomics Viewer (IGV) browser. Purple marks signal indels. Although not apparent in the long reads, the short-read mapping is consistent with a missing G (marked with a red arrow) at the end of the third repeat in the WD40 domain of *het-d*. White reads have multiple mappings. Blue reads signal smaller than expected insert size given the distribution of the paired-end reads.

**Figure S6.**
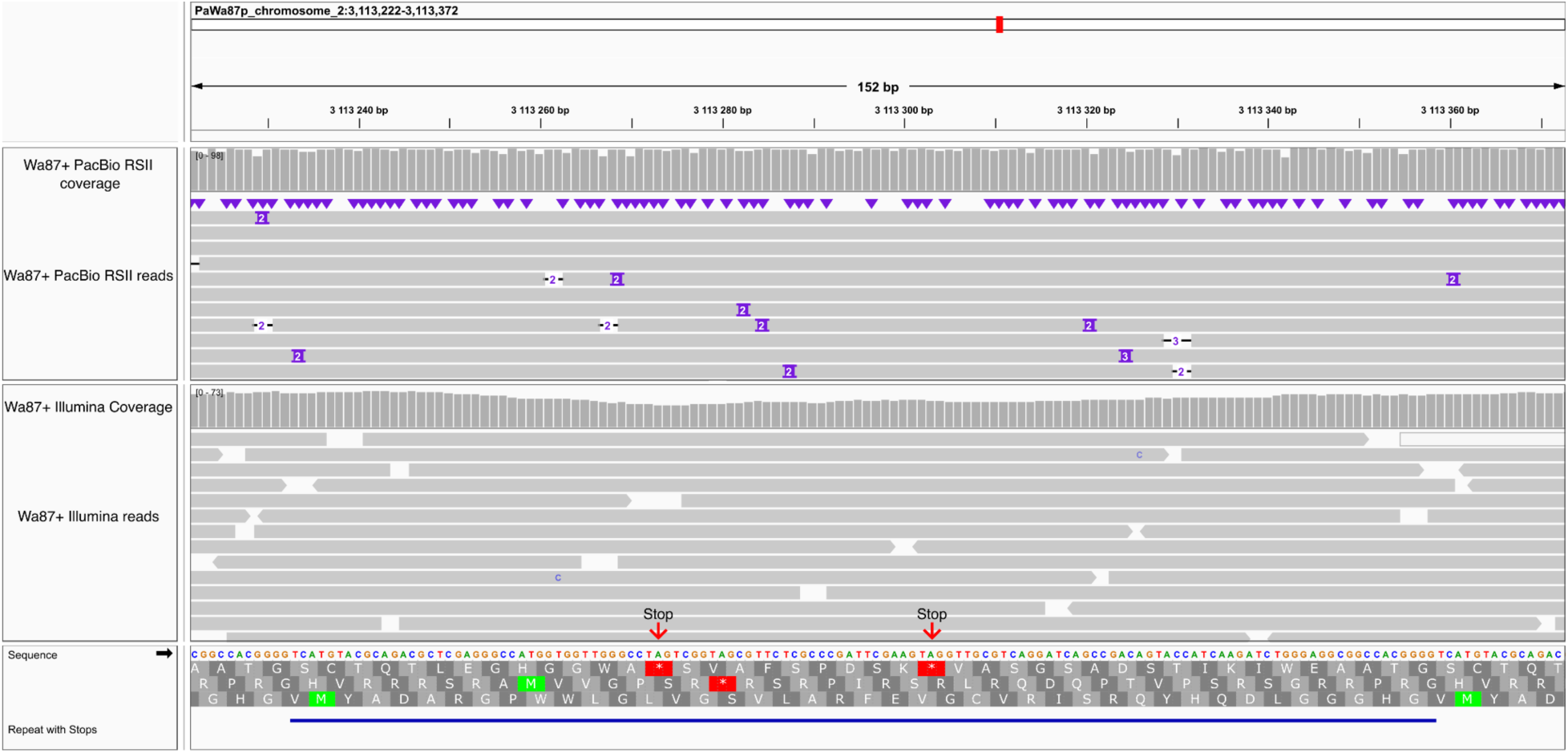
Short-and long-read mapping of the strain Wa87+ displayed in the Integrative Genomics Viewer (IGV) browser. Purple marks signal indels. The two stop codons found in the 6th repeat of the WD40 domain of *het-d* are marked with red arrows.

**Figure S7.**
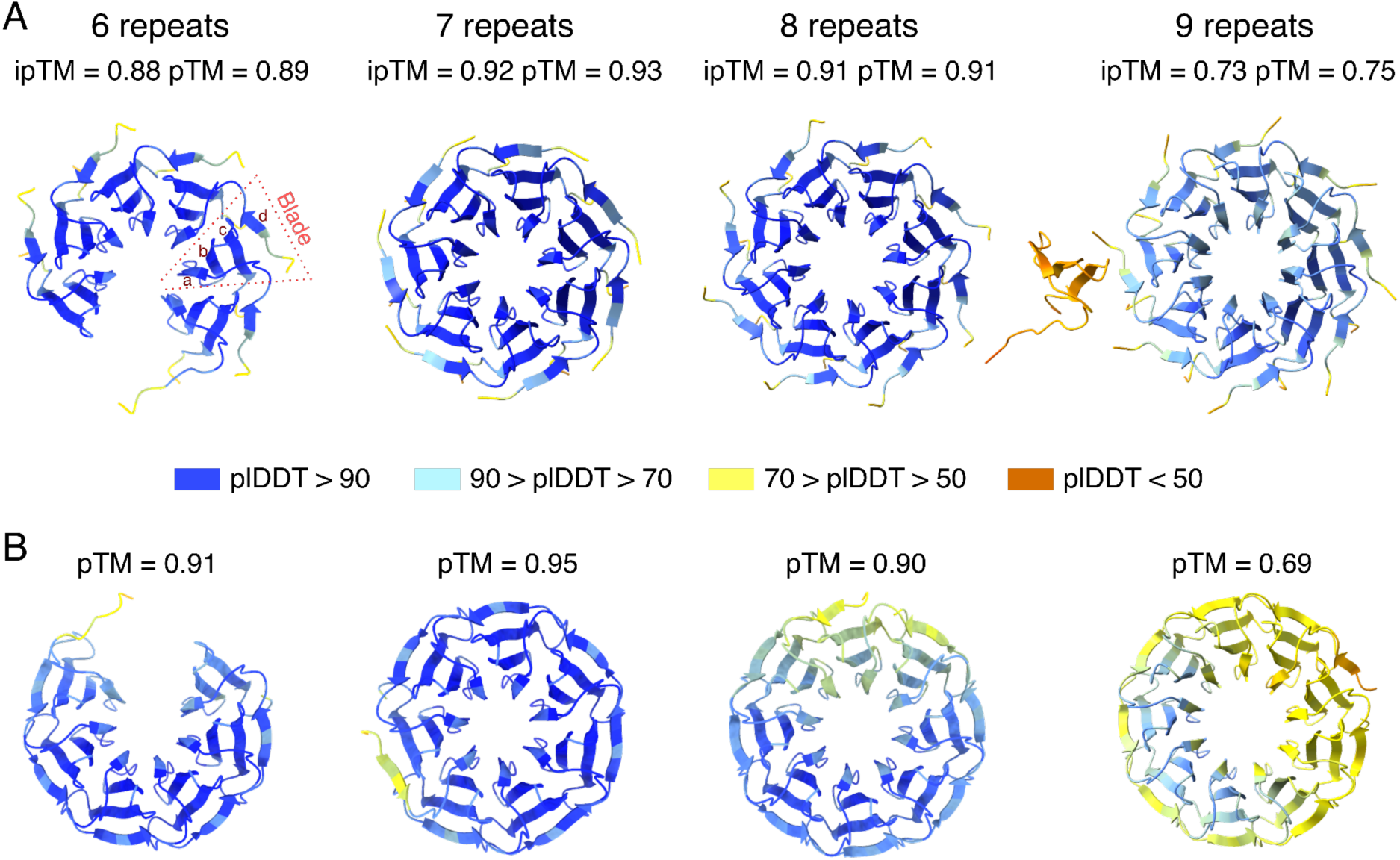
Ribbon diagrams of the β-propellers produced by AlphaFold 3 when different iterations of a HET-E2 repeat (second HIC repeat in the *E2*^C^ allele) are given. In (**A**) the individual repeat is input as multiple molecules to form a protein complex, while in (**B**) an artificial sequence with a given number of identical repeats folds into a single structure. The pLDDT score has a 0-100 scale where a higher value indicates higher confidence. The predicted template modeling (pTM) score and the interface predicted template modeling (ipTM) score have a scale from 0 to 1 and measure the accuracy of the entire structure (a score of 1 is best). The ipTM score in particular measures the accuracy of the relative positions of the subunits in the protein complex (in this case the blade monomers). An individual blade is formed by a *d* β-sheet from one repeat and the *a*, *b*, and *c* β-sheets of the next repeat, as highlighted in the first diagram of (**A**).

**Figure S8.**
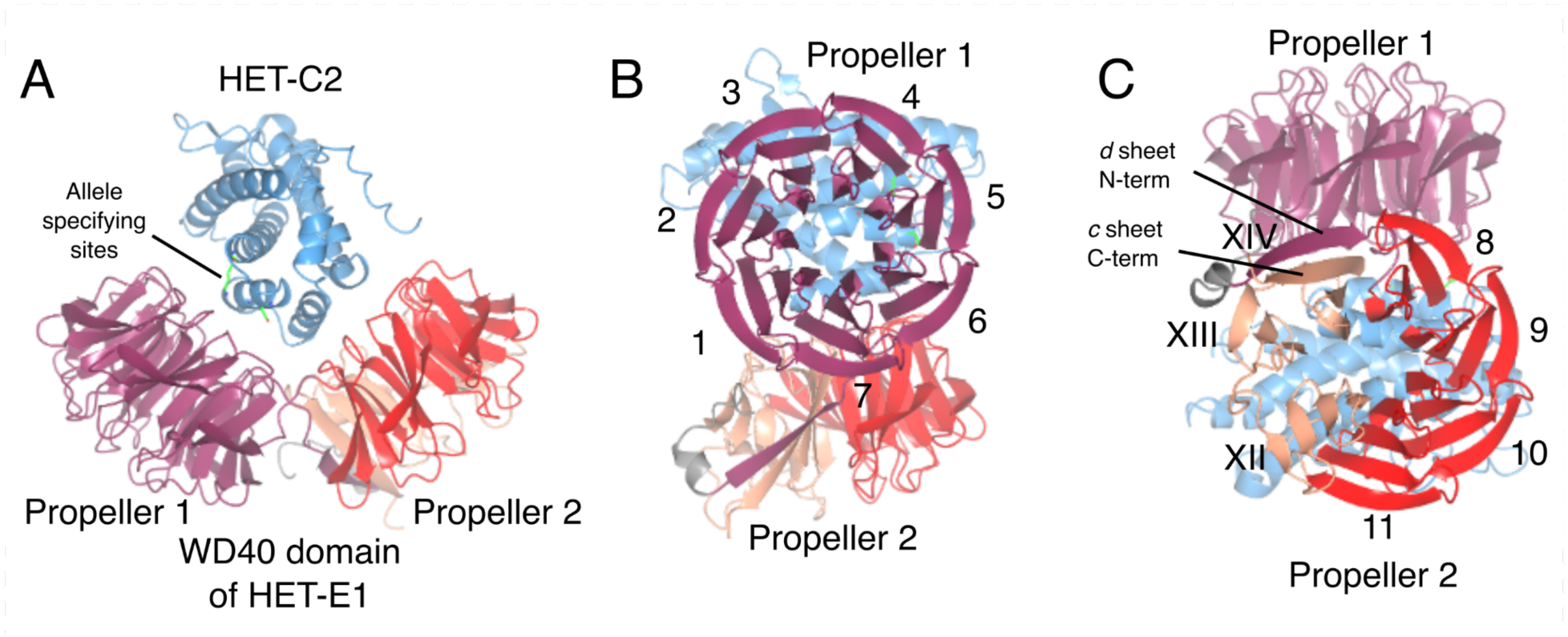
Ribbon diagrams of the WD40 domain of HET-E1^A^ interacting with HET-C2 as produced by AlphaFold 3. (**A**) The WD40 domain folds into two β-propellers with HET-C2 clamped in between. The first propeller is neatly assembled from seven HIC repeats (**B**), while the second propeller is produced from the remaining four HIC repeats, as well as three cryptic repeats (in Roman numerals) formed by the C-terminus of the protein and a *d* β-sheet on the N-terminus (**C**). The sites known to determine allele specificity are highlighted in HET-C2 with a stick representation (green).

